# Rainwater-driven transport of matter and microbes from phyllosphere to soil in a temperate beech forest

**DOI:** 10.1101/2025.07.13.664590

**Authors:** Markus Krüger, Karin Potthast, Beate Michalzik, Alexander Tischer, Martina Herrmann

## Abstract

In forest ecosystems, the transport of organic matter including microbial cells by throughfall and stemflow represents a central linkage between the phyllosphere and soil. However, the extent and taxon-specificity of rainwater-mediated vertical transfer of microorganisms, and its relevance for matter fluxes is sparsely understood. We investigated the rainwater-driven transport of bacteria and its contribution to matter transport in bulk rainfall, throughfall, stemflow and seepage in a temperate deciduous forest and compared the transported communities to those in phyllosphere and soil. Throughfall-mediated cell fluxes equaled those in seepage below the litter layer, pointing to a substantial above-ground translocation of bacteria. Bacterial cells contributed up to 11 % and 33 % to the exported total organic carbon and total nitrogen, respectively. Bacteria affiliated with *Pseudomonadaceae*, *Oxalobacteraceae*, *Hymenobacteraceae*, and *Sphingomonadaceae* were differentially transmitted from the canopy with increased abundances in throughfall and stemflow compared to bulk rain. A small fraction of shared taxa pointed to the import, proliferation and mobilization of selected bacterial taxa between neighboring ecohydrological compartments. The genetic potential of the transported bacterial communities was strongly linked to organic matter degradation particularly of carbohydrates and proteins. Our findings extend our understanding of rainwater-mediated vertical matter transport in forest ecosystems and highlight the contribution of microbial cells to this transport especially in above-ground compartments. Strong shifts in community composition and abundance of the transported taxa between throughfall, stemflow and soil seepage are linked to differences in source communities but may also point to different mechanisms of cell detachment across compartments.

**Highlights:**

- Rainwater flow paths are important carriers for organic matter and microorganisms.
- Throughfall and stemflow contained high abundances of microbial cells.
- Bacteria contributed up to 11 % and 33 % to exported carbon and nitrogen.
- Bacterial taxa were selectively mobilized by throughfall, stemflow and seepage.

## 1. Introduction

Rainwater represents an important carrier of various particles including organic matter and microorganisms across long distances (Hu et al., 2018). Such wet deposition may harbor tens of millions of bacteria m^−2^ d^−1^ (Reche et al., 2018) and other microorganisms such as archaea, fungi and protists (Despres et al., 2012; Sanchez-Parra et al., 2021), and thus plays a crucial role in microbial dispersal within ecosystems. Microbes from atmospheric sources may contribute directly to the microbiome of the plant phyllosphere (Ponette-Gonzalez et al., 2020), which exhibits a global surface area of 6.4 × 10^8^ km^2^ as microbial habitat (Vorholt, 2012). During rain events, huge amounts of rainwater can pass the forest canopy and emerge as throughfall and stemflow water fluxes (Levia et al., 2011). Throughfall represents that part of precipitation which falls through the canopy, partially along leaf surfaces, while stemflow is defined as the flow path of rainwater which runs down the stem (Sadeghi et al., 2020). In temperate forests, both water flow paths can transport a substantial amount of organic matter ranging between 0.2 – 200 kg C ha^−1^ y^−1^ (Ponette-Gonzalez et al., 2020; Van Stan et al., 2017) and bacteria in the range of 1.5 × 10^16^ cells ha^−1^ yr^−1^ (Bittar et al., 2018). As consequence, the forest canopy may act as a considerable source of microorganisms, influencing the bacterial composition of rainwater as it is transported down to the forest floor (Sadeghi et al., 2020; Van Stan et al., 2017). A certain proportion of this rainwater further percolates through the soil as seepage, containing mobile organic matter including microorganisms (Dibbern et al., 2014; Herrmann et al., 2023; Kruger et al., 2021), overall resulting in above-ground and below-ground microbial transport patterns .

In the phyllosphere, bacteria perform a variety of matter transformation processes such as carbon- and nitrogen fixation, nitrification and methanol oxidation (Andrews and Harris, 2000; Guerrieri et al., 2020). Methanotrophic bacteria such as *Methylomonas* and *Methylobacter, Flavobacterium* and *Novosphingobium* are common members of the microbiome on tree leaf and bark surfaces (Jeffrey et al., 2021; Vorholt, 2012). In addition, microbial extracellular enzymes which degrade hydrocarbons and secondary plant metabolites can be exported from the phyllosphere by throughfall and stemflow (Mori et al., 2019), contributing to exoenzymatic matter degradation in the soil. In contrast to litter fall in autumn, which acts as a periodic mass transfer of organic matter, microorganisms, and their degradation capacity to the forest floor, throughfall and stemflow represent season-independent, high-frequency episodic flow events from the phyllosphere, ensuring connectivity between above-ground and below-ground compartments and associated matter transfer throughout the year.

Assuming a certain fraction of the microbial cells remains viable during transport, rainwater delivers air- and phyllosphere-derived bacteria which constitute an essential source of new genetic material and functional potential to the sink communities. In fact, previous work demonstrated that some bacteria are selectively washed off from leaves and stems and get enriched along the water flow paths (Guerrieri et al., 2020; Teachey et al., 2018). Stemflow creates funneling of water along with matter and microbes which are rapidly transported along roots into the soil (Carlyle-Moses et al., 2020). In addition, rainwater provides fresh particulate and biolabile dissolved carbon and nitrogen to the forest floor (Michalzik et al., 2001; Van Stan and Stubbins, 2018), which can affect soil nutrient pools (Van Stan et al., 2017) and soil microbial activity (Deng et al., 2017).

Previous studies investigated the microbial diversity and mass fluxes associated with aerosol particles (Despres et al., 2012), tree phyllosphere, throughfall and stemflow (Van Stan et al., 2020). However, research is scarce on the quantity and structure of microbial communities transported by rainwater, throughfall, and stemflow, on which fraction of the transported matter originates from microbial cells, and on how their functional potential differs across different compartments including soil seepage. Seepage-mediated transport of microbes through soils was previously studied at the Hainich Critical Zone Exploratory using lysimeters installed in a beech forest (Kruger et al., 2021; Herrmann et al., 2023). In this study, we used zero tension lysimeters along with bulk rainwater, throughfall and stemflow collectors (Levia et al., 2023; Potthast et al., 2017) to investigate the abundance and community composition of bacteria along neighboring ecohydrological compartments ranging from the tree canopy to the soil.

These compartments included rainwater, phyllosphere, throughfall, stemflow, seepage and soil. In addition, we predicted the functional potential of these communities and their contributions to matter fluxes. We aimed to (1) estimate bacterial fluxes and contributions to carbon and nitrogen transport in throughfall, stemflow and seepage, comparing an early and late growing season rain event; (2) characterize the bacterial composition of the respective compartments and their predicted functional potential, and (3) assess core and shared communities for each intersection of neighboring compartments to provide insights into the fraction of bacterial communities that is subject to throughfall-, stemflow- or seepage-mediated mobilization.

## 2. Materials and methods

### 2.1 Study site and sampling procedure

Sampling was conducted in spring and autumn on a managed deciduous temperate forest plot (N51°06’56.5’ E10°24’45.2) and on an open area site (N51°07’63.5’ E10°23’68.2’) in the Hainich-Dun area comprising the Hainich Critical Zone Exploratory, Germany (**Figure 1A**). Forest stands are dominated by European beech (*Fagus sylvatica* L.) with minor admixtures of sycamore maple (*Acer pseudoplatanus* L.) and ash (*Fraxinus excelsior* L.), a common forest composition in the Hainich-Dun area. A nearby, extensively managed grassland site (open area) was used for measurements of bulk precipitation (rainwater). The climate is characterized by annual precipitation of 600 mm and mean air temperature of 9°C (Potthast et al., 2017).

**Figure 1:**
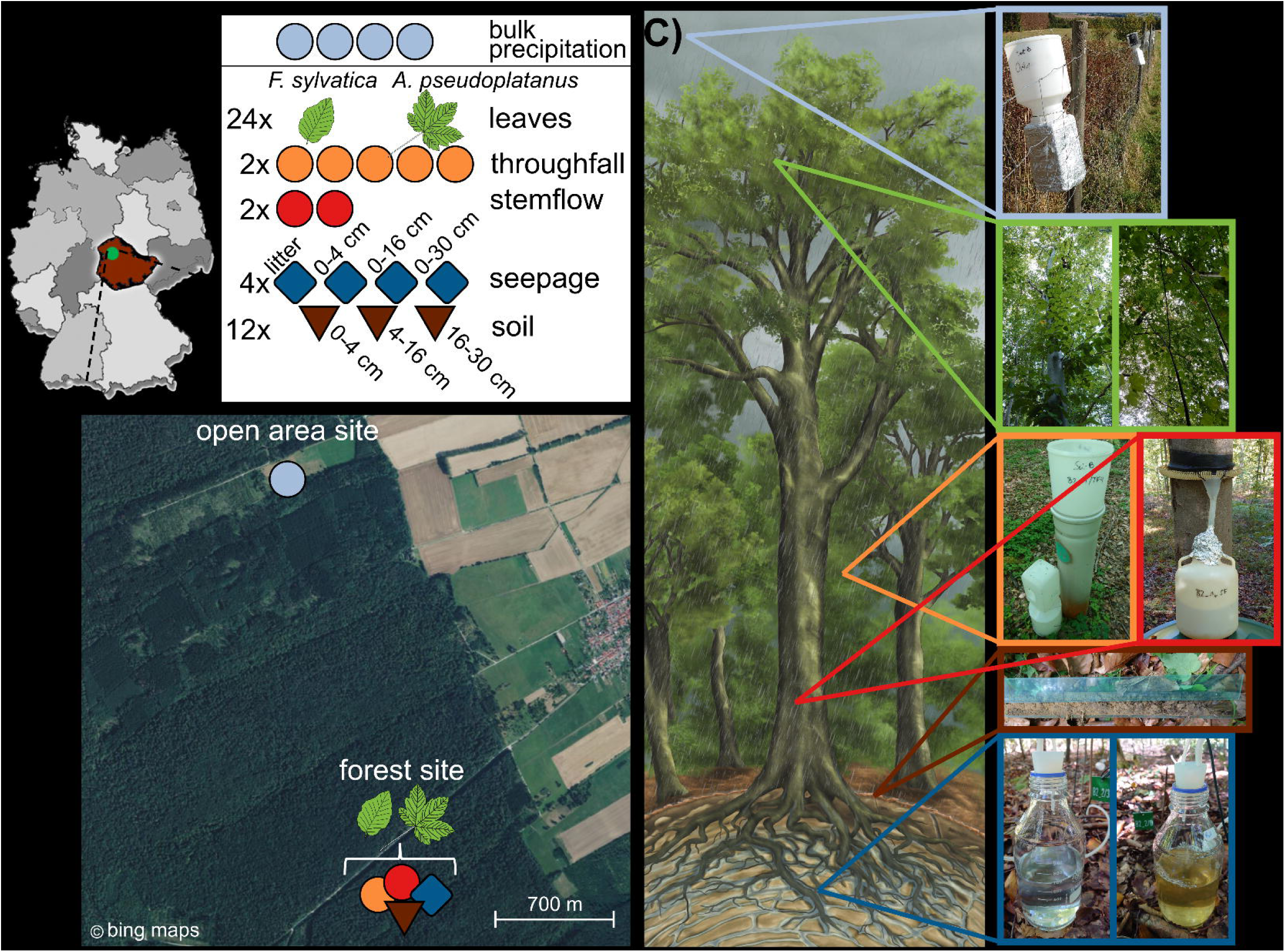
Study site and study design for sample collection at the Hainich Critical Zone Exploratory in western Thuringia, Germany. A) Sampling was conducted at an open area-site for rainwater representative of above-canopy precipitation and at a mixed beech forest site for phyllosphere, throughfall, stemflow, and seepage and soil sampling at different depths. B) Number of samples for each compartment during one sampling event. Sampling took place in May and September 2020. Soil was only collected in May. C) Conceptional representation of samples spanning from the tree canopy down to the soil, and photographs of the two tree species, sample collectors and a typical soil core.

Ten throughfall and four rainwater collectors with a sampling area of 314 cm^2^ at approximately 1.2 m height were deployed at a forest and an open area plot 2 km apart, respectively (**Figure 1B**). Stemflow collars were installed at two tree individuals of each *F. sylvatica* and *A. pseudoplatanus* at 1.3 m height with a basal area in breast height between 346 to 740 cm^2^. Rainwater, throughfall and stemflow plastic containers were cleaned and surface-disinfected with H2O2 prior to exposure and were placed in the field 1 to 2 days before two major rain events in May (expressed as early growing season) and September 2020 (expressed as late growing season), respectively. All three collector types were equipped with a 2 mm PE mash to avoid the introduction of large particles or insects during the collection period. Seepage was obtained from 16 zero-tension lysimeters with an area of 284 cm^2^ accessing soil leachate which percolated the soil through the litter, and the upper 4 cm, 16 cm and 30 cm (Kruger et al., 2021). Details on lysimeter properties and installation were described by Potthast et al. (2017). Residual solution from the lysimeters was emptied before the rain events to obtain only freshly percolated soil leachate. Throughfall, stemflow and seepage solutions at the forest site and bulk rainwater at the pasture site were sampled on 13^th^ May 2020 and 28^th^ September 2020 after rain events of 12 mm falling within 24 h and 19 mm falling within 48 h, respectively (**Figure 1C**). Triplicate leaf samples of the phyllosphere from two individuals of *F. sylvatica* and *A. pseudoplatanus* at low canopy height of 4 to 5 m were clipped off with ethanol cleaned scissors and stored in autoclaved plastic container after each rain event. In addition, soil samples as reference to the mobilized microbial communities were collected in triplicates close to the lysimeters at 0 to 4 cm, 4 to 16 cm and 16 to 30 cm depths in May 2020. All samples were stored in cooling boxes at approximately 8°C during transport and were immediately processed at the laboratory within 2 h. Rainwater, throughfall, stemflow, and seepage samples were weighed to determine the volume and 10 to 100 ml was used for determination of pH, electrical conductivity, carbon, nitrogen, and elemental analysis. For molecular analyses, water samples were filtered through 0.2 µm polyethersulfone membrane filters (Supor, Pall Corporation, USA) and stored at -80°C. Filtrate was stored at -20°C for determination of inorganic nitrogen. Phyllosphere samples in plastic bottles were treated with 1 min of mild sonification in 250 ml NaCl buffer (0.15 M NaCl + 0.1 % Tween 20) to detach microorganisms from leaf surfaces, followed by 20 min shaking at 100 rpm (Herrmann et al., 2021). Resulting suspensions were filtered through 0.2 µm pore size filters and stored at -80°C until further processing. The remaining leaves were dried at 50°C for 1 week to measure the dry weight. Soil samples for molecular analysis were stored at -80°C while those for determination of dry weight and pH were stored at 4°C.

### 2.2 Physicochemical analyses and calculation of matter fluxes

Rainwater, throughfall, stemflow, and seepage samples were analyzed for total organic carbon (TOC), inorganic carbon (IC) and total nitrogen (TN) as well as dissolved organic carbon (DOC) and total dissolved nitrogen (TDN) concentrations (see Dataset 1). Analysis involved filtration through <0.45 µm prewashed membrane filters (cellulose acetate, Sartorius, Gottingen, Germany) for dissolved elements and catalytic thermal oxidation and NDIR/chemiluminescence detection of both non-filtrated and filtrated samples using a TOC-VCPN/TNM-1 (Shimadzu, Duisburg, Germany) (Potthast et al., 2017). Particulate organic carbon (POC) and total particulate nitrogen (TPN) were calculated by subtracting the dissolved fractions from the total concentrations. Inorganic nitrogen (IN), mainly consisting of NH ^+^ and NO ^−^ was determined by standard colorimetric approaches (Scheiner, 1974; Seelmeyer, 1954).

Fluxes of TOC, TN, IN, and IC per h of rain events were calculated from the concentration of the respective parameters in the water samples (rainwater, throughfall, stemflow, seepage) multiplied by the respective sample volume and the collector area and were expressed in mg C or N m^−2^ h^−1^ as shown in Eq. 1 where *Cs* is the measured analyte concentration of a given sample, *Vs* represents the collected water volume, *Ac* is the area of the particular sample collector and *t* correspond to the rainfall period in h.

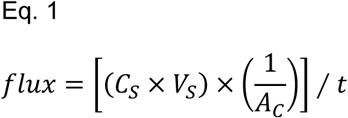

Stemflow fluxes were calculated for the individual basal area of each tree specimen and scaled up to the tree stand area in m^2^ per ha from both species.

Electrical conductivity and pH were determined using a WTW pH / Cond meter (pH Cond 3320 with SenTix81 and TetraCond, Weilheim, Germany) (Potthast et al., 2017). The soil water content was determined by drying the samples over 48 h at 105°C. Soil pH was determined from slurries prepared using sterile deionized water (ratio 1:2.5) and measured with a pH glass electrode (Kruger et al., 2021).

### 2.3 DNA extraction, quantitative PCR, and amplicon sequencing

Microbial biomass from a filter half (leaves, rainwater, throughfall, stemflow, seepage) and from 0.25 g of soil of each sample were extracted using the DNeasy PowerSoil kit (Qiagen, Germany) following the manufacturer’s protocol and stored at -20°C. Bacterial and archaeal 16S rRNA gene abundances were quantified with a CFX96 real-time PCR thermocycler (Bio-Rad, USA) using primer pair Bakt_0341F/Bakt_799R (Klindworth et al., 2013, Chelius and Triplett, 2001) which discriminates against chloroplast-derived 16S rRNA genes, and Arch806F/Arch958R (DeLong, 1992; Takai and Horikoshi, 2000), respectively. Details on PCR and cycling conditions are described elsewhere (Herrmann et al., 2021; Kruger et al., 2021). Plasmids containing cloned bacterial and archaeal 16S rRNA gene fragments were used to construct qPCR standards, which were linear across the concentration range of 50 to 5 ×10^8^ copies per reaction. Gene abundances in rain, throughfall, stemflow and seepage were calculated as per L, while gene abundance on leaf surfaces and soil was expressed as genes per g of dry weight (see Dataset 1).

Library construction for 16S rRNA gene targeted amplicon sequencing assessing the bacterial community composition across samples was conducted with the same primer pair as for qPCR, targeting the V3-V4 region of the bacterial 16S rRNA gene. Reactions were performed with 1 ng DNA template per sample in 15 µl volume reactions under previously reported conditions (Herrmann et al., 2021). Subsequent steps are described in Kruger et al. (2021), and sequencing was performed on an Illumina MiSeq platform using v3 chemistry.

### 2.4 Sequence processing

PCR primer sequences were trimmed from raw reads using Cutadapt (Martin, 2011) and reads were further processed with the R package DADA2 1.22.0 following the standard sequence analysis pipeline for 16S rRNA genes (Callahan et al., 2016). Trimmed reads were filtered and denoised using the filterAndTrim function under the following conditions: (truncLen = c(270, 240), maxEE = c(2,3), truncQ = 2, maxN = 0). The filtered reads of each sample were dereplicated, Amplicon Sequencing Variants (ASVs) were deduced for forward and reverse reads separately, and then the forward and reverse reads were merged at overlapping positions. Chimeric sequences were removed and taxonomic assignment of amplicon sequence variants (ASV) was performed using the SILVA taxonomy reference database release version 138 (Quast et al., 2013). Subsequent sequence analysis was conducted using the package phyloseq (McMurdie and Holmes, 2013) and is further described in the Supplementary Methods.

### 2.5 Prediction of microbial functions

To approximate the functional potential of the microbial communities across the different compartments, PICRUSt2 (Phylogenetic Investigation of Communities by Reconstruction of Unobserved States) analysis was applied to predict microbial functions based on the obtained amplicon sequencing data using v2.3.0 beta following the standard procedure for 16S rRNA (Douglas et al., 2020). ASV sequences were first aligned and placed into a reference tree and ASVs were corrected for multiple 16S rRNA genes by the 16S rRNA operon number of closely related representatives. Gene families for each ASV were predicted based on KEGG orthologs (KO) as identifiers for metabolic pathways (Kanehisa et al., 2023). The PICRUSt2 output of 7,523 KOs was subset for KO abundance of enzymes involved in the following pathways: carbon fixation, nitrogen fixation, methane oxidation, carbon degradation and protein hydrolysis (see Dataset 4). Metagenomic predictions of the selected pathways were based on the relative abundance of the KO IDs across each sample. To assess the proportion of the bacterial community associated with the selected metabolic functions across samples, we used the relative abundance of each ASV from amplicon sequencing data for which a corresponding KO was predicted (see Dataset 4).

### 2.6 Determination of absolute cell abundances and cell fluxes

Studies have shown that combining amplicon sequencing of marker genes and qPCR can be used as an accurate method to infer cell numbers similar to direct cell determination by flow cytometry (Bittar et al., 2018) and to enhance bacterial community profiling (Jian et al., 2020). The quantity of exported cells was calculated using the 16S rRNA qPCR-based gene abundances per liter and the amplicon sequencing reads on family-level (see Dataset 2). To account for diverging operon numbers of the 16S rRNA gene across the 303 observed bacterial families, gene abundances were corrected using the number of operons of the respective family-level taxa obtained from the ribosomal RNA operon database (https://rrndb.umms.med.umich.edu/) (Stoddard et al., 2015) (see Dataset 3). The estimated total cell abundance per liter (*Tabs*) for each sample of rainwater, throughfall, stemflow and seepage was determined based on the relative read abundance from each taxon divided by the family-specific operon numbers and multiplied by the total 16S rRNA gene abundance. This calculation is expressed in Eq. 2 where *Sabs* represents the total gene abundance of a given sample, *Trel* is the relative abundance of a taxon and *Tn* represents the number of 16S rRNA operons for a taxon.

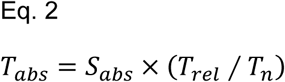

Bacterial cell fluxes per m^2^ h^−1^ were calculated as described for TOC and TN using the estimated absolute cell abundance per total sample volume and collector area for each sample, divided by the duration of the respective rain event (see Dataset 1).

### 2.7 Estimation of bacterial contributions to TOC and TN

The fraction of bacterial cell carbon and nitrogen contributing to TOC and TN in rainwater, throughfall, stemflow and seepage was estimated as previously described (Herrmann et al., 2023; Kruger et al., 2021). Briefly, a mean carbon content of 39 fg per cell and mean nitrogen content of 12 fg per cell was assumed as estimated under carbon-limited and nitrogen-limited conditions (Vrede et al., 2002), respectively. The contribution of bacterial cell carbon and nitrogen to TOC and TN, respectively, was estimated by first multiplying the operon number corrected absolute bacterial abundances as well as the TOC/ TN content per liter by the sample volume. The estimated number of bacterial cells was multiplied by the assumed carbon and nitrogen content per cell. The resulting estimates of microbial cell carbon and nitrogen were then set in relation to the total organic carbon (TOC) and total nitrogen (TN) in the corresponding samples to estimate the bacterial contribution (Dataset 3).

### 2.8 Statistical analysis

All statistical tests were performed using R version 4.1.2 (R Core Team, 2020) unless otherwise described. Normal distribution of gene abundances and element fluxes were tested with Shapiro-Wilk test. Non-parametric Kruskal-Wallis rank sum test and Dunn’s multiple comparison post-hoc test with R package FSA (Ogle et al., 2020) was used to test data for significant differences between the compartments. Nonmetric multidimensional scaling using Bray-Curtis distance matrix was performed to test dissimilarity of the bacterial community between the compartments, the growing seasons and the tree species. More details on statistical methods and the R packages used are provided as Supplementary Methods.

## 3. Results

### 3.1 Carbon and nitrogen fluxes across compartments

The export of total organic carbon (TOC) ranged from 0.1 mg C m^−2^ h^−1^ in stemflow to 45.1 mg C m^−2^ h^−1^ in seepage right below the litter layer and was significantly different between compartments (*p* < 0.001) (**Figure 2A**). Inorganic carbon (IC) fluxes were on average 2 to 4 times lower across all compartments (**Figure S 1**). Total nitrogen (TN) flux varied from as low as 0.01 mg N m^−2^ h^−1^ in stemflow to a maximum of 29.3 mg N m^−2^ h^−1^ in seepage collected after percolation through the upper 16 cm of the soil profile (**Figure 2A**). C:N ratios showed significantly higher values in throughfall and stemflow (*p* < 0.001) compared to rainwater and seepage (**Figure S 1**). Ammonium flux decreased from rainwater, throughfall and stemflow to the seepage flow path while fluxes of nitrate were highest in seepage (**Figure S 1**).

**Figure 2:**
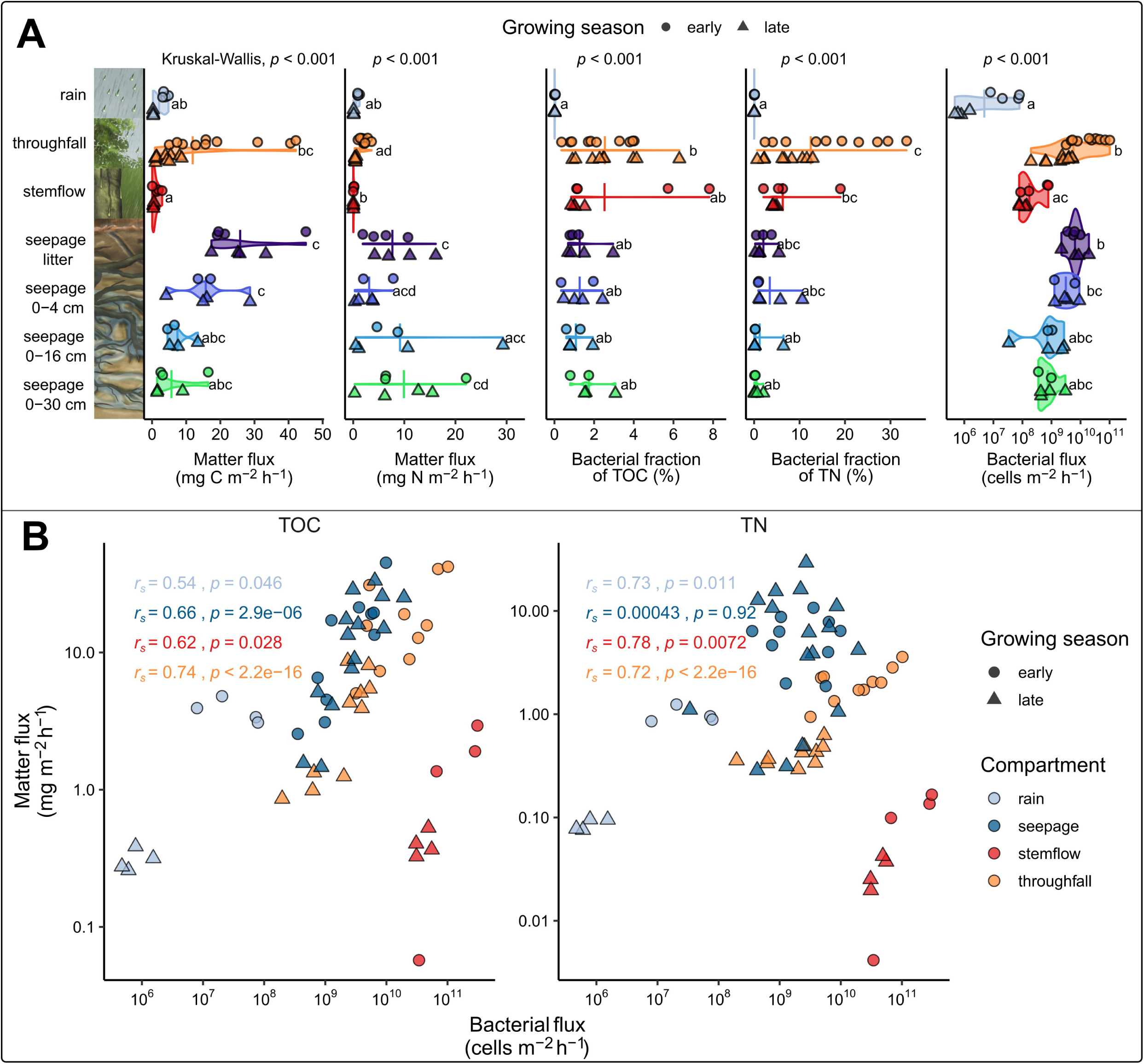
Bacterial transport across the different compartments, and contributions and correlations to transported carbon and nitrogen. **A)** Fluxes of TOC and TN expressed per m^2^ h^−1^, bacterial contributions to TOC and TN content, and fluxes of bacterial cells per m^2^ h^−1^ across the different compartments during an early and late growing season rain event. Each symbol represents a single sample, and the crossbar depicts the average for each compartment. Significant differences are shown in letter code tested by post hoc Dunn’s multiple comparing (*p* < 0.05). **B)** Spearman’s rank correlation (*rs*) between bacterial cell flux per m^2^ h^−1^ and TOC and TN of water fluxes per m^2^ h^−1^ in May and September.

### 3.2 Microbial gene abundances, estimated cell fluxes, and contributions to C and N transport across compartments

Bacterial 16S rRNA gene abundances per L (**Figure S 2**A) differed significantly between the different compartments (*p* < 0.001) and were highest in throughfall and stemflow. Rainwater exhibited on average three orders of magnitude lower gene abundances. In seepage, bacterial 16S rRNA gene abundances decreased with increasing collection depth from below the litter layer down to 30 cm soil depth. Archaeal 16S rRNA gene abundances were consistently two to three orders of magnitude lower than bacterial 16S rRNA gene abundances and exhibited an opposing depth-dependent trend in seepage.

We further estimated bacterial cell abundances per g and L based on qPCR data and amplicon sequencing data, applying family-level corrections for multiple 16S rRNA gene operons per cell (**Table S 1**). Estimated bacterial abundances on leaf surfaces ranged from 1.4 × 10^5^ to 5.4 × 10^8^ cells g^−1^ dry leaf material while soil bacterial abundances ranged from 9.6 × 10^8^ to 7.8 × 10^9^ cells g^−1^ dry soil. In rainwater, estimated bacterial cell abundance ranged from 1.2 × 10^6^ to 1.6 × 10^8^ cells L^−1^ while throughfall and stemflow cell abundances were about 100 times higher, ranging from 5.8 × 10^8^ to 3.1 × 10^11^ cells L^−1^ and 7.9 × 10^9^ to 2.3 × 10^11^ cells L^−1^, respectively. Seepage bacterial abundance ranged from 3.8 × 10^8^ to 3.9 × 10^10^ cells L^−1^ with a decreasing trend with soil depth (*p* < 0.01).

Based on estimated cell abundances, we also determined the flux of bacterial cells per m^2^ and h transported by rain, throughfall, stemflow and seepage during both sampling events. Across all samples and the two sampling time points, cell fluxes ranged from lowest values in rainwater with 4.7 × 10^5^ cells m^−2^ h^−1^, 1.9 × 10^10^ cells m^−2^ h^−1^ by seepage from the litter layer to highest export of 1.0 × 10^11^ cells m^−2^ h^−1^ by throughfall with significant differences between compartments (*p* < 0.001) (**Figure 2A**). On average, cell fluxes via throughfall and litter layer were three orders of magnitude higher than in rainwater. Seepage-mediated cell fluxes of seepage collected below 4 cm soil depth were one order of magnitude lower than in throughfall. No differences in estimated cell fluxes were observed between early and late growing season for a given compartment.

Fluxes of TOC and TN by rainwater, throughfall and seepage were positively correlated with the estimated total bacterial cell flux across all compartments and sampling months (*p* < 0.05) (**Figure 2B**), underlining the strong relationship between microbial transport dynamics and matter transport. Estimation of the bacterial contribution to matter transport indicated a significant bacterial contribution across the different water flow pathways (*p* < 0.001) with bacterial cells contributing between 0.006 % of TOC and 0.007 % TN in rainwater, and up to 11.3 % TOC and 33.6 % TN in throughfall **(Figure 2A)**.

### 3.3 Bacterial community composition from phyllosphere to soil

The ASV richness and alpha diversity (Shannon index) differed significantly between all compartments (Kruskal-Wallis, *p* < 0.001) (**Figure 3A**). The highest number of different bacterial taxa was observed in seepage, followed by stemflow, soil, leaf surface and throughfall samples, while rainwater samples were the least diverse. Leaf, seepage, and soil communities were on average more diverse than communities from rain, throughfall and stemflow samples (*p* < 0.05). Alpha diversity and species richness of throughfall and stemflow communities were lower during early growing season compared to late growing season (*p* < 0.05) while this effect was not observed for the other compartments.

**Figure 3:**
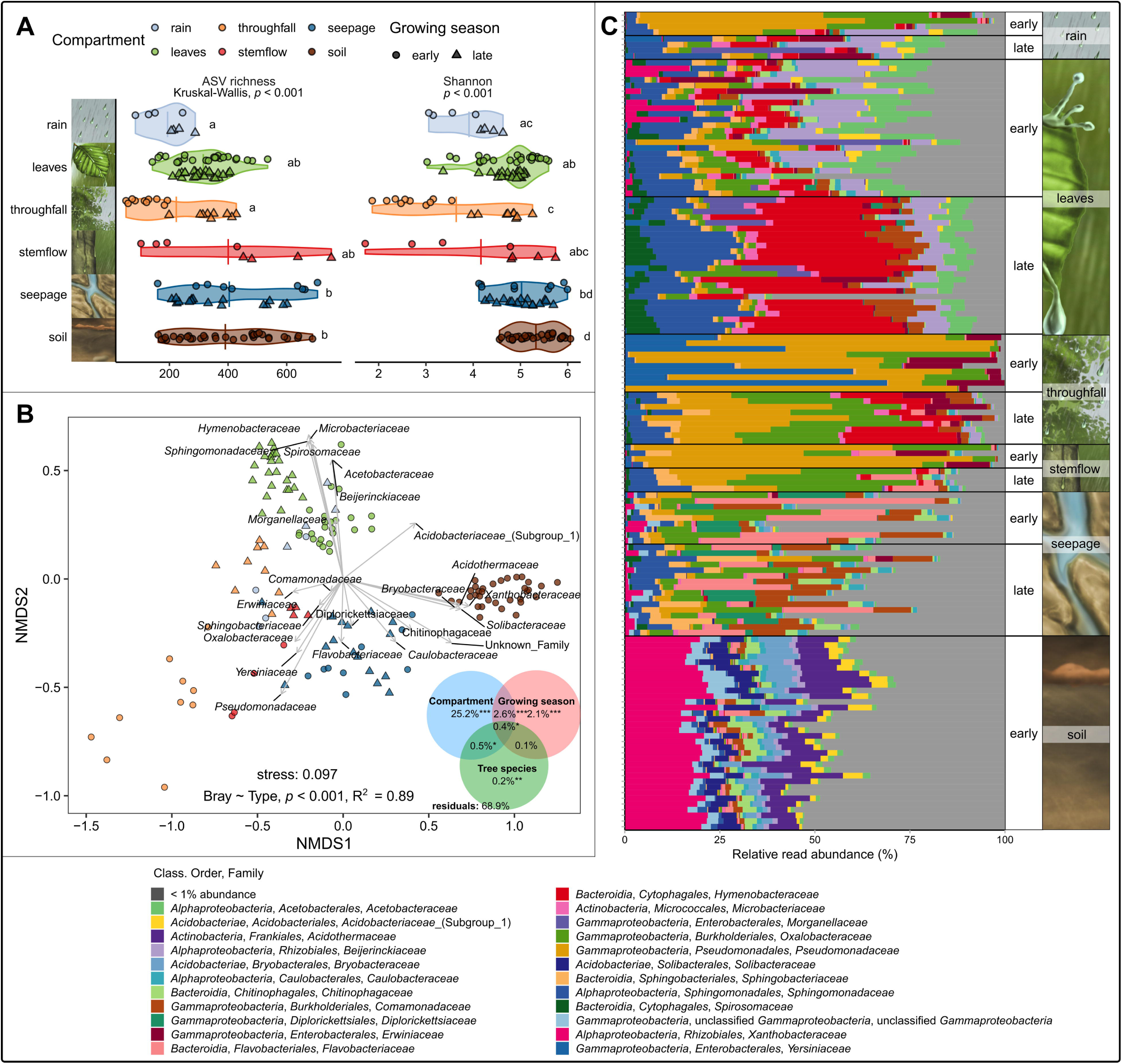
Bacterial communities across rainwater (n = 8), leaves (n = 48), throughfall (n = 20), stemflow (n = 8), seepage (n = 25) and soil (n = 34) samples. **A)** Observed ASV richness (left panel) and Shannon index (right panel). Single data points are shown as dots for samples from May and triangles from September (crossbar represents mean of values). Letter code indicates significant differences between compartments and indexes after Dunn’s post hoc test (*p* < 0.05). **B)** Distribution of most abundant bacterial families depicted as NMDS based on Bray-Curtis dissimilarities across all samples. Results of PERMANOVA (9999 permutations) are displayed as Venn diagram, asterisks depict confidence interval at *p* < 0.001 for each accounting factor. **C)** Bacterial community composition of most abundant bacterial families with more than 1 % absolute read abundance per sampling point. Single stacked bars show the relative read abundance of 16S rRNA gene sequences on family level.

Beta diversity analysis of bacterial communities revealed a clear differentiation based on the sampled compartment (ANOSIM, R^2^ = 0.52, *p* < 0.001) with phyllosphere, seepage and soil communities being distinct, while rainwater, throughfall and stemflow communities appeared very variable and were not clearly separated (**Figure 3B**). The compartment was the strongest driver for community differentiation, accounting for 25.2 % of the total community variation (PERMANOVA, *p* < 0.001) (**Figure 3B**). The sampling month contributed 2.1 % of the variation. The influence of tree species (European beech versus sycamore maple) explained only less than 1 % of the variation and was not included as an influencing factor in further analysis.

On family-level, 23 bacterial taxa comprised each more than 1 % of the total community (**Figure S 2**B), which exhibited distinct distribution patterns across rain, throughfall, stemflow and seepage (ANOSIM: R^2^ = 0.54, *p* < 0.001). Gammaproteobacteria, Alphaproteobacteria, Bacteroidia, Actinobacteria and Acidobacteriae accounted for at least 90 % of all sequence reads across all samples (**Figure 3C**). Across rainwater, throughfall and stemflow samples, *Pseudomonadaceae* accounted for a large proportion of between 24 – 42 % together with common plant pathogens such as *Erwiniaceae* and *Yersiniaceae*, particularly during the early growing season. In phyllosphere samples, *Hymenobacteraceae* (32 mean ± 7 % standard deviation) dominated at the late growing season sampling point together with *Sphingomonadaceae* (16 ± 7 %) while *Beijerinckiaceae* (13 ± 6 %) were more abundant at the early growing season. In contrast to rainwater, throughfall and stemflow, Gammaproteobacteria were less abundant in phyllosphere samples and were here mostly represented by *Oxalobacteraceae* (5 ± 3 %) and *Comamonadaceae* (6 ± 4 %) in the early and late growing season samples, respectively. In soil communities, Alphaproteobacteria accounted on average for 24 ± 3 % with *Xanthobacteraceae* being the dominant family (19 ± 3 %). Seepage communities were dominated by *Flavobacteriaceae, Oxalobacteraceae* and *Comamonadaceae*.

The taxonomic composition derived from estimated cell abundances confirmed the patterns revealed by relative read abundances (**Figure S 2**C). Analysis based on absolute abundance revealed that most of rainwater-derived bacterial families, including *Pseudomonadaceae*, *Yersiniaceae* and *Sphingomonadaceae*, were two to five orders of magnitude more abundant in throughfall and stemflow. In contrast to those communities, the bacteria in seepage depicted a two to three orders of magnitude increase in cell abundance of taxa such as *Xanthobacteraceae*, *Spirosomaceae* and *Diplorickettsiaceae,* which showed also high relative abundances in soil communities.

### 3.4 Predicted functions of bacterial communities

The prediction of the bacterial genetic repertoire for selected biogeochemical transformations unveiled distinct patterns across the compartments (**Figure S 3**, ANOSIM: R^2^ = 0.42, *p* < 0.001). The predicted relative abundance of KOs showed that most pathways had higher prevalence towards soil and leaf communities except for protein hydrolysis, which was more prevalent in bacteria transported by seepage.

The functional metagenomic prediction was based on the presence and absence of functional modules of each pathway (**Figure 4A**). Hydrolytic enzymes showed high relative abundances of methionyl aminopeptidase (0.083 %) and leucyl aminopeptidase (0.062 %) across the samples. Potential for the degradation of carbon sources was also highly prevalent across the samples depicted by beta-N-acetylhexosaminidase (0.046 %), hexosaminidase (0.033 %), beta-glucosidase (0.029 %), chitinase (0.022 %) and alpha-amylase (0.017 %). Potential for carbon fixation was mostly associated with the Calvin–Benson–Bassham pathway, represented by *cbbS* and *cbbL* genes with an average relative abundance of 0.002 % and 0.006 %. Nitrogenases as primary enzymes for nitrogen fixation were prevalent in soil (0.0037 %), phyllosphere (0.0022 %) and seepage communities (0.0018 %). Methane oxidation was primary related to methanol dehydrogenase (0.0014 %) across all samples, with the highest relative fraction in phyllosphere (0.0037 %) and rainwater communities (0.0024 %).

**Figure 4:**
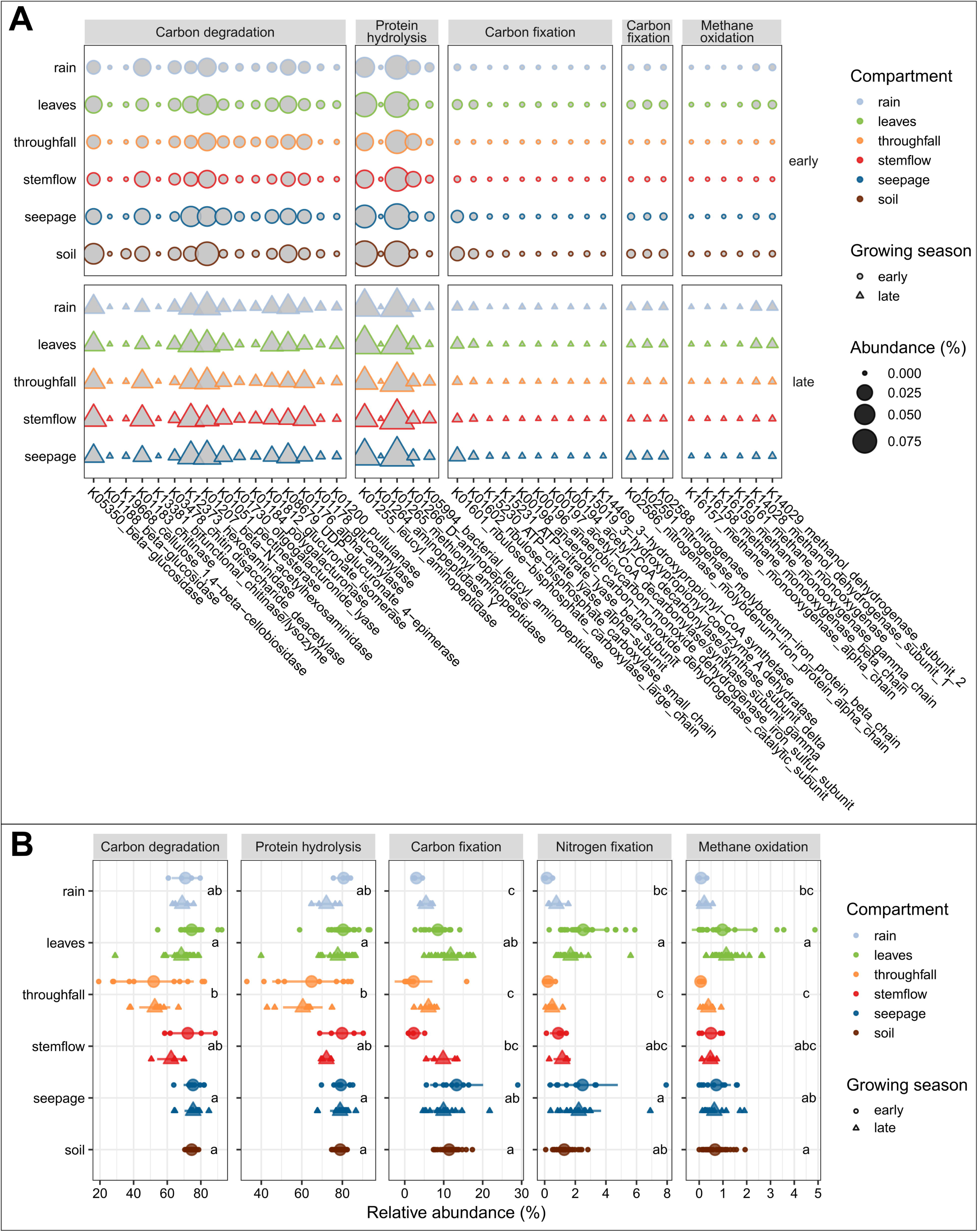
Prediction of metabolic functions of bacterial communities across the different compartments and the two time points. **A)** Abundance of KEGG orthologues (KO) corresponding to genes for enzymes of selected metabolic pathways. The size of the bubbles depicts the mean relative KO abundance of each compartment. **B)** Proportions of bacterial communities employing the predicted KO. Small dots depict the mean relative abundance of bacterial classes per sample and the bigger dots represent the mean relative abundance for a given compartment. Letter code displays significant differences between compartments within each pathway based on post hoc Dunn’s multiple comparing (*p* < 0.05).

A similar pattern of the functional profiles across the compartments was observed based on relative abundance of bacterial classes for which predicted KO were present (**Figure 4B**). The prevalence of bacteria which showed functional potential differed significantly between the compartments for each pathway (*p* < 0.05). The fraction of bacterial communities showing capability of utilizing enzymes involved in various carbon and protein decomposition processes ranged, on average, from 52 % in throughfall to 75 % in seepage and 60 % in throughfall to 80 % in leaf samples, respectively. Carbon fixation made up by a fraction of 2 % to 13 % of the bacterial community with lowest potential in throughfall and stemflow, and highest potential in seepage samples. Pathways related to biological nitrogen fixation and methane oxidation showed the highest prevalence of 2.5 % and 1.1 %, respectively, in leaf communities, while rain and throughfall communities showed functional potential for both pathways of less than 1 %.

### 3.5 Association of bacterial taxa with rainwater- and seepage-driven transport

Only a small proportion of 0.19 % (23 ASVs from a total of 12191 ASVs without singletons) of the bacterial community was shared across all compartments. These shared ASVs were especially abundant in the communities that were transported by rainwater, throughfall or stemflow, with ASVs related to *Pseudomonas*, *Massilia* and *Sphingomonas* accounting for up to 33% of sequence reads in the respective communities (Figure S 4). The major fraction of 62 % of all ASVs observed in this study were exclusively present in seepage, while less than 10 % of the total ASVs were only found in either rain, throughfall or stemflow communities. Stemflow communities shared 5.9 % of the ASVs with rainwater, 10.7 % with throughfall, and 6.9 % with seepage.

We defined bacterial core taxa based on occurrence in at least 70 % of all samples and tracked the increase or decline of single core ASVs across the different compartments, along with an assessment of their potential transport by throughfall, stemflow, and seepage. Core taxa which constituted a large community fraction on leaves included members of the *Hymenobacteraceae*, *Acetobacteraceae*, *Bejierinckiaceae* and *Sphingomonadaceae*. ASVs affiliated with the genera *Hymenobacter*, *Acidiphilium*, *Methylorubrum* and *Sphingomonas* accounted for up to 1 % relative abundance of the total community on leaves but were less abundant in rainwater. Some ASVs classified as *Sphingomonas* were also observed in throughfall and stemflow (**Figure 5**, **Figure S4**). Among the phyllosphere community, *Acidiphilium*, *Methylorubrum* and *Sphingomonas* were also observed in rainwater. In contrast, *Pseudomonadaceae* and *Oxalobacteraceae* including the genera *Pseudomonas* and *Massilia*, respectively, were abundant in rainwater, contributed on average 41% ± 24% and 14% ±11% of the communities in throughfall and 42% ± 30% and 13% ±8% in stemflow, but were less abundant in seepage with 3.3% ± 3.7% and 3.6% ± 2.8% and on leaves (3% ± 5.2% and 3.3% ± 2.8%, respectively).

**Figure 5:**
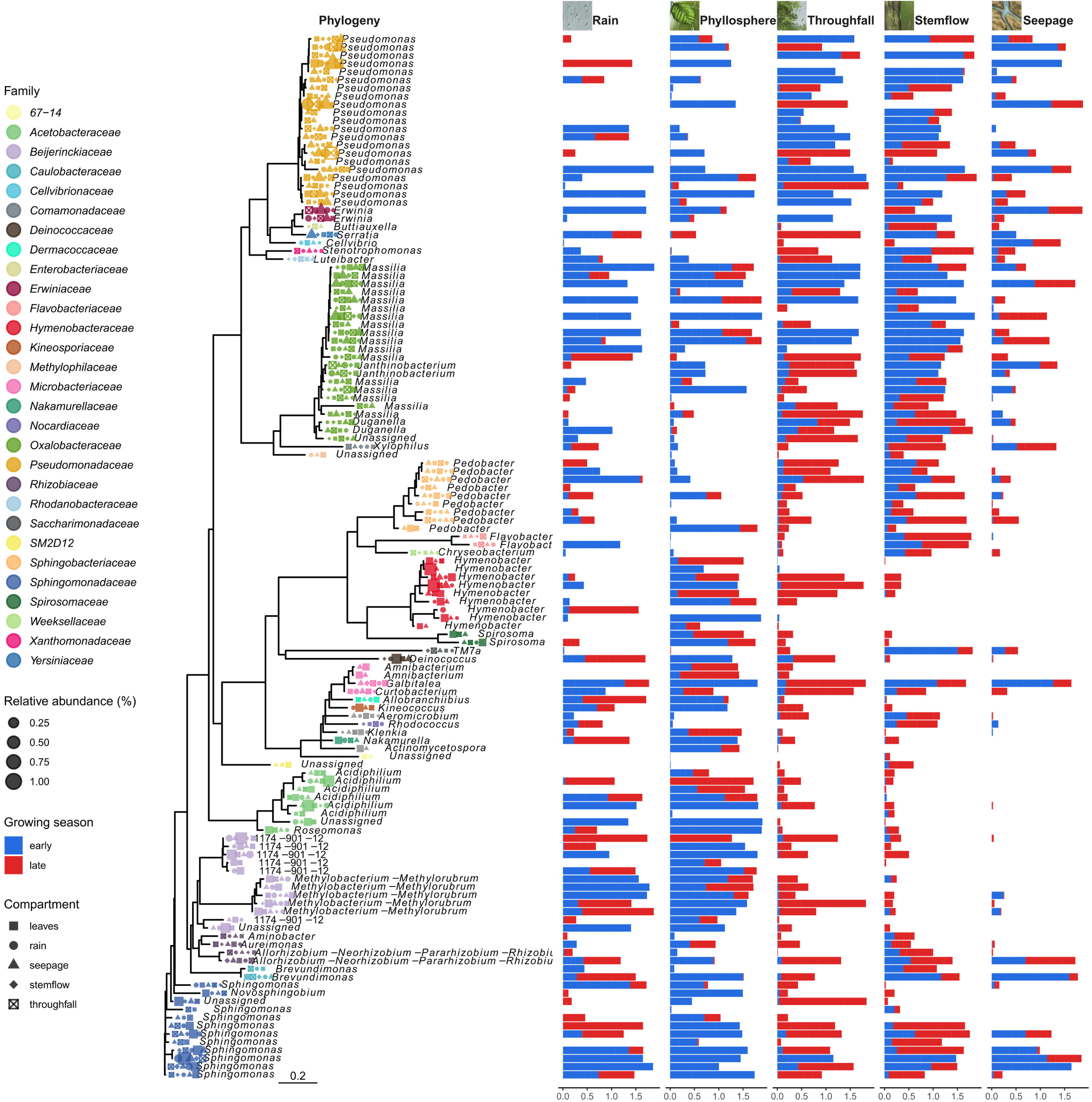
Phylogeny and relative abundance of bacterial core taxa across rainwater, phyllosphere, throughfall, stemflow and seepage samples during both sampling time points. The tree is based on a neighbor-joining model of a 16S rRNA gene alignment. The color and shape of each node show the presence of an ASV on family-level and on genus-level, respectively, at a given compartment. The size of nodes depicts the mean relative abundance of each ASV across the compartments. Stacked bars display the mean relative abundance of each ASV across all samples from both sampling points of rain, phyllosphere, throughfall, stemflow and seepage communities.

In addition to the preferential association of core ASVs with a specific compartment, we determined the differential abundance of genus-level taxa which showed either the tendency to stick on leaf surfaces or to be washed off by throughfall and stemflow (**Figure S 5**), indicating their potential to become subject to rainwater-mediated transport. Members of *Pseudomonadaceae*, *Yersiniaceae* and *Oxalobacteraceae* exhibited high relative abundances in throughfall and stemflow while *Sphingomonadaceae* and *Hymenobacteraceae* showed stronger association with the phyllosphere. Instead, members of *Xanthobacteraceae* and *Flavobacteriaceae* showed higher relative abundances in seepage compared to throughfall and stemflow (**Figure S 6**), and *Flavobacteriaceae* and *Sphingomonadaceae* formed a larger fraction of the seepage community compared to soil. LEFSe analysis confirmed the specific association of those bacterial taxa with either the tree phyllosphere or rainwater-, throughfall- or stemflow-associated mobilized fractions (**Figure S 7**).

## 4. Discussion

Precipitation events transmit a vast number of organic particles, including microbes, representing a major environmental agent connecting the atmosphere with the phyllosphere, pedosphere and lithosphere (Ponette-Gonzalez et al., 2020; Van Stan et al., 2020). The transported matter and microbial cells can potentially impact microbial communities and nutrient cycling along the water flow path from the plant phyllosphere to soil (Van Stan and Stubbins, 2018; Van Stan et al., 2020). However, the extent of rainwater-driven microbial mobilization within the forest phyllosphere and from bark surfaces, and its contribution to carbon and nitrogen fluxes have so far remained unknown. Here, we assessed the diversity and abundance of bacteria along this flow path, starting from intercepting rain via tree canopy-derived throughfall and stemflow to soil seepage in a temperate forest area at an early and late stage of the growing season.

Our results clearly demonstrate that canopy-derived throughfall and stemflow microbial communities are quantitatively important carriers of organic matter to the forest floor, contributing up to 11 and 34 % to total carbon and nitrogen fluxes, respectively. Thus, they play a central role in the throughfall- and stemflow-mediated provision of fresh organic carbon and nitrogen to the forest floor (Van Stan et al., 2021). Throughfall and stemflow obtained in this study from a beech forest during a two-days rain event were highly enriched with bacteria with 10 to 1000 times higher estimated cell numbers than previously reported from a subtropical oak-cedar forest (Bittar et al., 2018). Given the fact that throughfall and stemflow samplers were directly sampled at the end of a three-days exposure period at overall low ambient temperatures between 0-20°C, we can exclude that massive growth of bacterial cells in the samplers may have led to a strong overestimation of these bacterial loads. Despite on average similar cell concentrations L^−1^ in throughfall and stemflow solutions, stemflow-mediated cell fluxes normalized to tree stand area and h were two orders of magnitude lower than in throughfall, which is typically also observed for TOC and TN fluxes when comparing these two water flow paths (Van Stan and Stubbins, 2018). Throughfall cell fluxes were comparable to those from seepage collected below the litter layer, indicating a substantial above-ground translocation of bacteria to soil.

In contrast to the high estimated proportions of bacterial derived carbon and nitrogen in throughfall and stemflow, bacterial contributions to TOC and TN were three times lower in seepage solutions, matching with previously reported values from the same lysimeters (Kruger et al., 2021) and tension-controlled lysimeters from the same field site (Herrmann et al., 2023). Total estimated cell fluxes were even ten times lower in seepage below 16 cm compared to throughfall. Apparently, a large fraction of the cells mobilized by throughfall and stemflow get retained in the soil, which acts as a strong filter for this above-ground transport. Turnover processes in the soil result in carbon and nitrogen export primarily as dissolved compounds with a lower contribution of mobilized microbial cells. During rain events, throughfall and stemflow mediated inputs might supplement the nutrient pool of the forest litter and soil and could additionally affect microbial nutrient cycling capabilities by introducing active microorganisms with their respective biodegradation capabilities or exoenzymes from upstream communities. Pathway predictions indeed indicated that bacterial communities from throughfall, stemflow and seepage exhibited strong potential for hydrolytic degradation of carbohydrates and proteins. However, since PICRUSt2 only predicts metagenomic functions from marker gene abundance and phylogenetic placement to reference sequences (Douglas et al., 2020), speculations about the full degradation potential need to be treated with caution.

The type of canopy-derived transport agent (throughfall versus stemflow) was the strongest effector for bacterial community similarity/ dissimilarity, suggesting that the composition of the phyllosphere versus bark-associated source communities and potential differences in microbial detachment mechanisms between these two habitats could underlie the observed community differences. Phyllosphere communities were more similar to rain, throughfall and stemflow communities than to seepage and soil, suggesting an exchange of microbes between phyllosphere, throughfall and stemflow in a taxon-specific manner. Rainwater can potentially support the colonization but also the wash-off of epiphytic microbes from leaf surfaces (Guerrieri et al., 2020). The observed fraction of 10 % of shared ASVs between throughfall and stemflow communities suggested that runoff from the canopy exported a basic, selective fraction of bacteria independent of the phyllosphere location, which was further modulated by exchange with the specific colonizers of either leaves or bark (Guerrieri et al., 2020; Teachey et al., 2018). Similarly, Hudson et al. (2023) demonstrated that the bacterial community composition in stemflow was distinct from that found on tree bark. Bark-water interactions during rainfall may lead to the detachment of loosely attached microbes and enable their accumulation further down as infiltration source in soil (Carlyle-Moses et al., 2020).

Weather conditions and tree species were identified as important factors shaping throughfall bacterial communities in a mediterranean holm oak forest (Guerrieri et al., 2020) and a subtropical Southern live oak forest (Teachey et al., 2018) with Alphaproteobacteria, Gammaproteobacteria and Bacteroidia representing main bacterial classes. Following the above-ground flow path at our study site, phyllosphere-inhabiting *Pseudomonadaceae*, *Erwiniaceae*, *Oxalobacteraceae* and *Sphingobacteriaceae* may partially have originated from rainfall-mediated atmospheric inputs of early leave colonizers and exhibited 10 to 100 times higher relative abundances on leaf surfaces compared to rainwater. In turn, they also appeared to be selectively transported by throughfall and stemflow, indicating an overall high mobilization potential of these taxa. Our results further demonstrated a substantial difference in the microbial communities of the above-ground flow paths – rainwater, throughfall and stemflow - when comparing an early and late time point of the growing season. During May sampling, *Pseudomonadaceae* accounted on average for over 50 % of the total community in bulk rain, throughfall and stemflow, indicating that at this time point, only a single taxonomic group contributed substantially to matter transport and potential carbon degradation capabilities. *Pseudomonas* species act as common pathogens and epiphytic plant commensals in the phyllosphere (Vorholt, 2012). However, they are also typically observed among microbes attached to plant pollen (Manirajan et al., 2018), which can be transmitted by dry deposition (Sanchez-Parra et al., 2021) and rainwater (Cho and Jang, 2014). Higher pollen bioaerosols or pollen deposits derived from the flowering of beech or nearby Norway spruce stands in May might have constituted a large microbial source of pollen-associated *Pseudomonadaceae* in rainwater, throughfall and stemflow (Guidone et al., 2021).

The observed distinction of bacterial communities in seepage versus throughfall and stemflow may not only be linked to differences in the bacterial source communities but also to differences in the general bacterial detachment mechanisms between above-ground and below-ground habitats. Preferential, taxon-specific microbial transport through soil by seepage-mediated translocation is mainly attributed to cellular characteristics such as cell size, shape and surface charge (i.e., zeta potential) (Dibbern et al., 2014; Herrmann et al., 2023). In the phyllosphere, the main factor controlling the preferential transfer from leaf surfaces to throughfall might be the strength of attachment to the surface rather than cell size. Members of the *Pseudomonadaceae* can increase local water availability of leaf surfaces by releasing surfactants, allowing proliferation (Koskella, 2020), which might also be a reason for preferential mobilization and detachment of members of this family by throughfall and stemflow. In addition, surfactants may also support the detachment of other leaf-associated microorganisms. Prevalent bacterial epiphytes from the phyllosphere beyond *Pseudomonadaceae* are *Beijerinckiaceae* and *Sphingomonadaceae* (Delmotte et al., 2009; Vorholt, 2012), which were also observed in throughfall and stemflow at our study site. Especially *Pseudomonadaceae* could also constitute an important source of extracellular exoenzymes performing hydrolytic degradation, affecting decomposition of organic matter (Mori et al., 2019).

Despite the high numbers of cells transported by throughfall and stemflow, it remains an open question whether cells remain viable throughout the whole vertical passage, and how many living microbes ultimately reach the soil. A seepage study using electron microscopy indicated that living microbes remained intact during soil passage across 60 cm depth by seepage transmission (Dibbern et al., 2014). However, it is unclear whether the throughfall- and stemflow-mediated cell transfer to the soil mainly contributes to inputs of organic carbon and nitrogen, or whether it adds to the functional potential and enzymatic capacity of the local soil microbial populations.

Our results further suggest that multiple taxa were able to colonize and thrive in all of the investigated ecosystem compartments from atmosphere to soil. Establishment and proliferation in the respective compartments probably ensure that the taxa can also be released again from these compartments and be transported by throughfall, stemflow or seepage. However, with our approach, we cannot distinguish between continuous cross-compartment transport of individual cells or discontinuous transport, resulting from establishment in each compartment, proliferation, and subsequent secondary mobilization of microbial taxa. Consequently, occurrence of the same ASVs across the vertical flow path does not necessarily indicate an uninterrupted flow of individual cells through the system but the ability of certain taxa to be continuously mobilized from one compartment and transported to the other, and to establish stable populations in the respective sink communities, from whereon cells can be mobilized again, resulting in the observed overlaps of microbial communities and their functional potential between neighboring ecohydrological compartments.

## 5. Conclusion

Our study clearly demonstrates that microorganisms transported by rainwater, throughfall, stemflow and seepage contribute substantially to carbon and nitrogen fluxes in a deciduous forest ecosystem. Contributions and cell fluxes were larger in above-ground compared to below-ground compartments, suggesting that a considerable fraction of mobilized cells was retained in the soil and not further transported. Throughfall- and stemflow-mediated flow paths act as substantial carriers of microbial biomass, carbon and nitrogen which might affect soil matter transformation by introducing a fresh source of nutrients and energy but also new genetic variability to the respective microbial sink communities. In contrast to periodic cell and nutrient inputs via litter fall, the throughfall- and stemflow-mediated vertical microbial transport to the forest floor takes place throughout the year, acting as central connection between phyllosphere and forest floor in terms of carbon and nitrogen inputs but also as contribution to the functional diversity of soil microbiomes. Our results further expand a previously observed taxon-specificity in seepage-mediated microbial mobilization in soil to throughfall and stemflow, presumably along with different mechanisms of microbial detachment across these flow paths.

## Data availability

Raw sequence files were deposited at the European Nucleotide Sequence Archive with project number PRJEB92466 under accessions ERS25231161 to ERS25231303.

## Acknowledgements

We thank Falko Gutmann, Stefan Riedel and Linda Gorniak for support of field and laboratory work, and Martin Stephan from the Otto Schott Institute for Materials Research at Friedrich Schiller University Jena for producing customized soil lysimeters. We are further grateful for the background graphics provided by Clemens Schulze.

## Author contributions: CRediT

**Markus Kruger**: Conceptualization, Formal analysis, Investigation, Methodology, Visualization, Writing – original draft, Writing – review and editing. **Karin Potthast**: Conceptualization, Methodology, Writing – review and editing. **Beate Michalzik**: Conceptualization, Writing – review and editing. **Alexander Tischer**: Conceptualization, Methodology, Writing – review and editing. **Martina Herrmann**: Funding acquisition, Conceptualization, Sample processing, Writing – review and editing.

