## Supplemental Material for "Rainwater-driven transport of matter and microbes from phyllosphere to soil in a temperate beech forest"

Markus Krüger[1], Karin Potthast[2,3], Beate Michalzik[2,3], Alexander Tischer[2,3,4], Martina Herrmann[1,3,5*]

[1]Institute of Biodiversity, Ecology and Evolution, Friedrich Schiller University Jena, Jena, Germany; [2]Department of Soil Science, Friedrich Schiller University Jena, Jena, Germany; [3]German Center for Integrative Biodiversity Research (iDiv) Halle-Jena-Leipzig, Leipzig, Germany, [4]ThüringenForst, Forestry Research and Competence Centre, 99867 Gotha, Germany, [5]Cluster of Excellence Balance of the Microverse, Friedrich Schiller University Jena, Jena, Germany,

### Supplemental Methods

#### Amplicon sequence analysis

After assignment of ASVs, sequences were extracted and aligned using DECIPHER (Wright, 2016), and a phylogenetic tree was constructed using phangorn (Schliep, 2011). In depth analysis of the bacterial community was performed in R using the packages phyloseq (McMurdie and Holmes, 2013), vegan (Oksanen et al., 2008) and tidyverse (Wickham et al., 2019) and their dependencies. Non-bacterial sequences assigned to chloroplasts, mitochondria, eukaryotes, and archaea were excluded from further analysis, resulting in 15659 unique ASVs and a median of 20,464 reads per sample. One seepage sample was excluded from analysis due to very low amounts of sequencing reads per sample. Normalization was done by rarefying all reads from each sample using the phyloseq function rarefy_even_depth at a minimum of 4426 sequencing reads per sample. Subsequently, ASVs with only 1 read were excluded from analysis resulting in a total of 12191 ASVs across all samples. Shannon and Chao alpha diversity indexes were calculated using the estimate_richness function in phyloseq.

#### Statistical analysis

All statistical tests were performed using R version 4.12 (R Core Team, 2020) unless otherwise described. Normal distribution of gene abundances and element fluxes were tested with Shapiro-Wilk test. Non-parametric Kruskal-Wallis rank sum test and Dunn's multiple comparison post-hoc test with R package FSA (Ogle et al., 2020) was used to test data for significant differences between the compartments. The significance of the observed differentiation was tested with ANalysis Of SIMilarity (ANOSIM) using the anosim2 function from vegan. Correlations between microbial gene abundance, TOC, TN and the sample volumes were assessed and visualized using ggpubr (Kassambara, 2023). Relationships between the absolute abundance of the most abundant 17 bacterial families and the matter content of water solution samples were determined by pairwise correlation using Spearman's rank coefficient and visualization using ComplexHeatmap (Gu et al., 2016) and its dependencies.

The number of shared ASV across the all samples was assessed using packages phyloseq, MicrobiotaProcess (Xu et al., 2023) and VennDiagram (Chen and Boutros, 2011). The prevalence of dominant ASV representing the core microbiome across the samples was determined using the metagMisc package including phyloseq, vegan, ALDEx2 (Fernandes et al., 2013), metagenomeSeq (Paulson et al., 2013) and DESeq2 (Love et al., 2014).

The shared core microbiome of leaf, rain, throughfall and stemflow was assessed by filtering ASV which were present in at least 70 % of the samples using the function phyloseq_filter_prevalence from metagMisc. The sequences of core AVSs from leaf, rain, throughfall and stemflow samples were extracted and placed in a phylogenetic tree using ggtree (Xu et al., 2022). Lastly, the relative abundances of those ASV during both growing seasons were plotted next to the tree as described in (Rolando et al., 2022).

The differential abundance of abundant ASV between phyllosphere and canopy-derived flow paths was assessed using the package DESeq2 (Love et al., 2014). Unrarefied sequence reads with > 1 sequence across all samples were subset for leaf, throughfall and stemflow compartments. The phyloseq object was converted to a DESeq2 data file and the ASV table was transformed using the median ratio method for differential abundance analysis.

The package microbiomeMarker (Cao et al., 2022) was utilized to investigate the bacterial community for biomarker ASV on genus level. Linear discriminant analysis (LDA) Effect Size (LEFSe) was performed across all samples using CPM normalization method with a *p* value of 0.05 and LDA cutoff of 4.

To assess the contributing factors accounting for the similarity/ dissimilarity of the bacterial community across compartments, a Permutational Multivariate Analysis of Variance Using Distance Matrices (PERMANOVA) was conducted using a Bray-Curtis distance matrix and the adonis2 function with 9999 permutations in R package vegan.

#### Figure S1

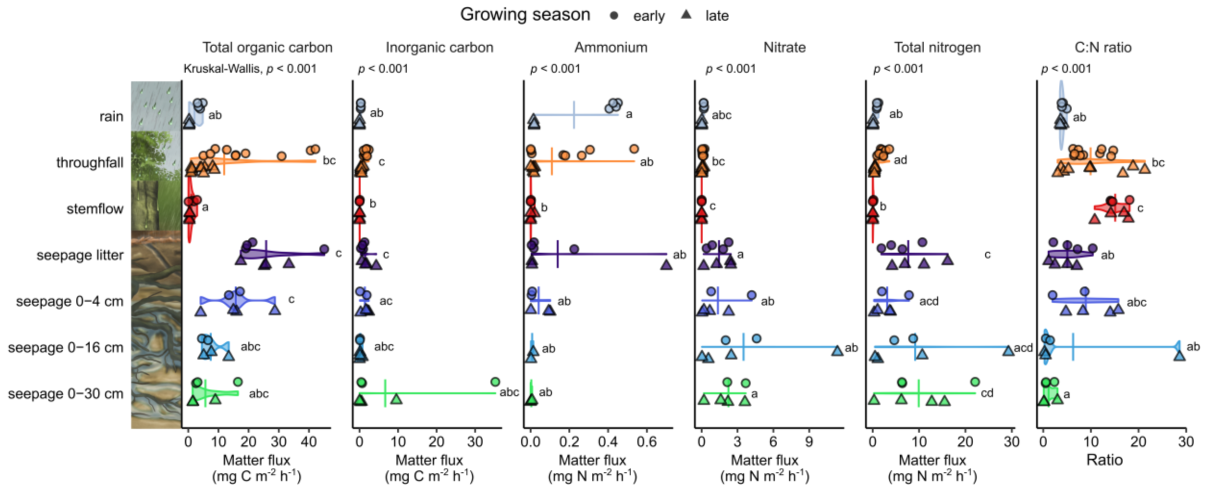


**Figure S 1:** Fluxes per $m^2$ and h of total organic carbon, inorganic carbon, ammonium-N, nitrate-N and total nitrogen, and C:N ratios (fluxes of TOC+IC/TN) from water solutions during early and late sampling point. Significant differences are shown in letter code tested by Dunn's multiple comparing test ($p < 0.05$).

### Figure S2

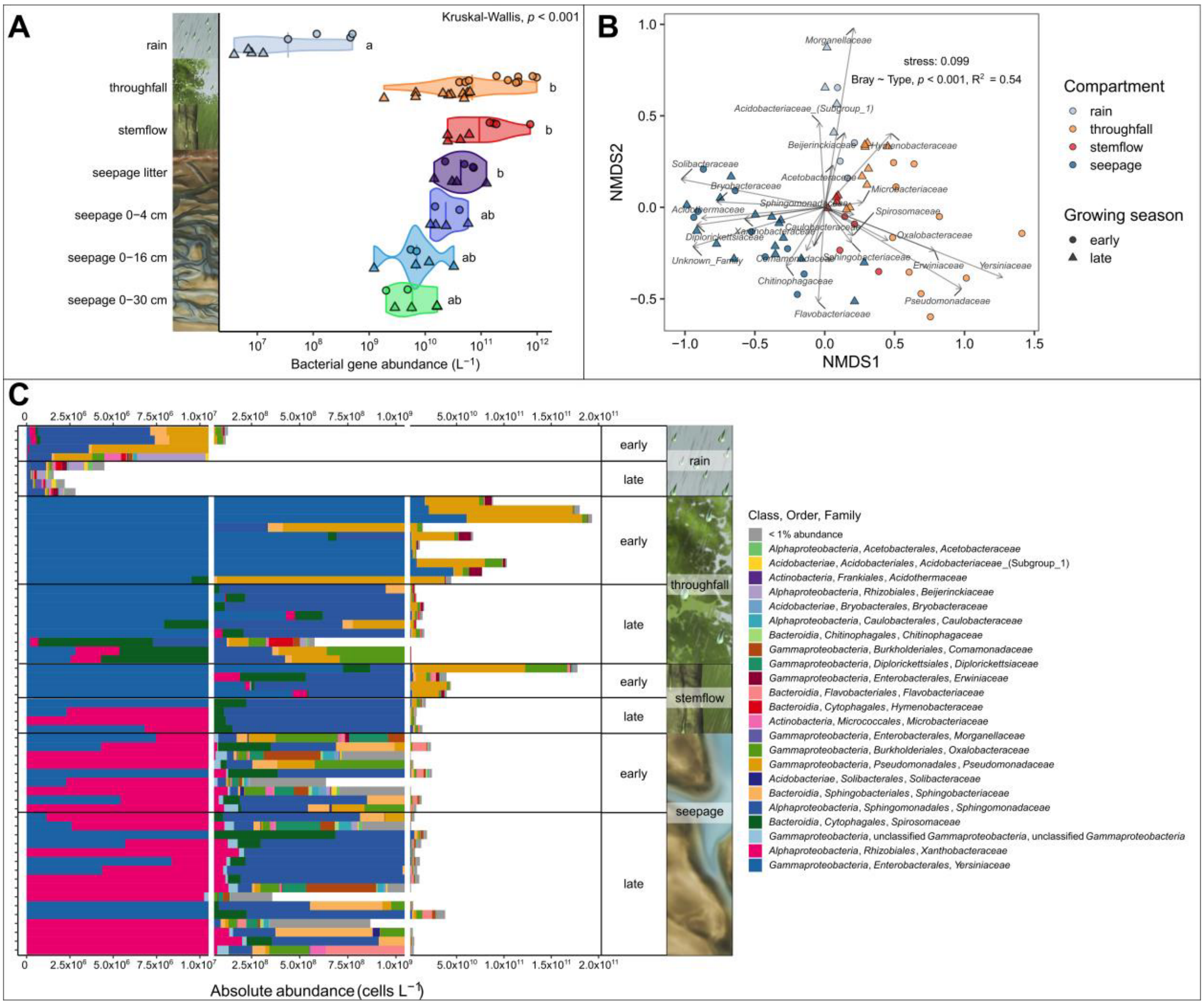


**Figure S 2:** Transport of bacteria by rainwater, throughfall, stemflow and seepage. **A)** Bacterial gene abundance per liter across the water flow paths. The circle (May) and triangle (September) symbols depict single samples and the crossbar depicts the average for each compartment. Significant differences are shown in letter code tested by Dunn's multiple comparing test ($p < 0.05$). **B)** Distribution of the total bacterial community across compartments and both sampling points using NMDS based on Bray-Curtis dissimilarity matrix of absolute cell abundance. Significant differences of the communities between compartments are shown as result of an ANOSIM test. **C)** Bacterial community composition of most abundant bacterial families and with less than 1 % absolute. Single stacked bars show the absolute cell abundance per liter derived from of the 16S rRNA genes obtained by qPCR and corrected by inferring the operon numbers on family-level.

**Figure S3**

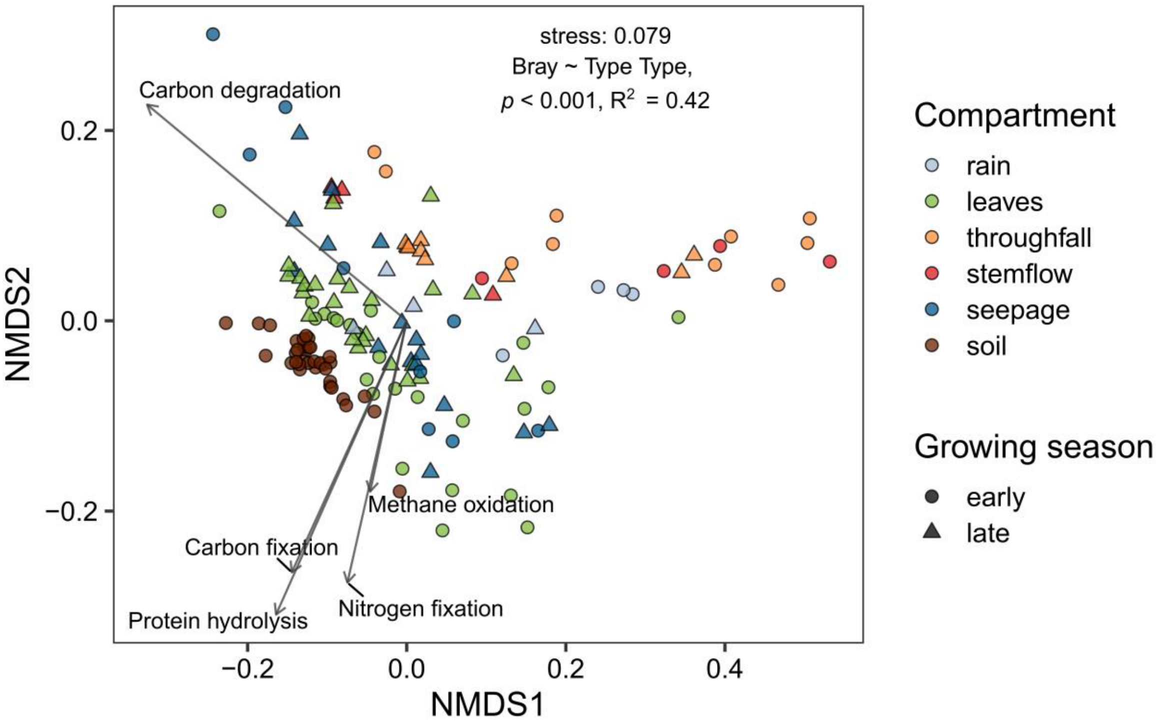


**Figure S 3:** Distribution of the predicated metabolic pathways across all compartments based on summarized KO abundance per compartment and sampling point shown as NMDS based on Bray-Curtis dissimilarity matrix.

**Figure S4**

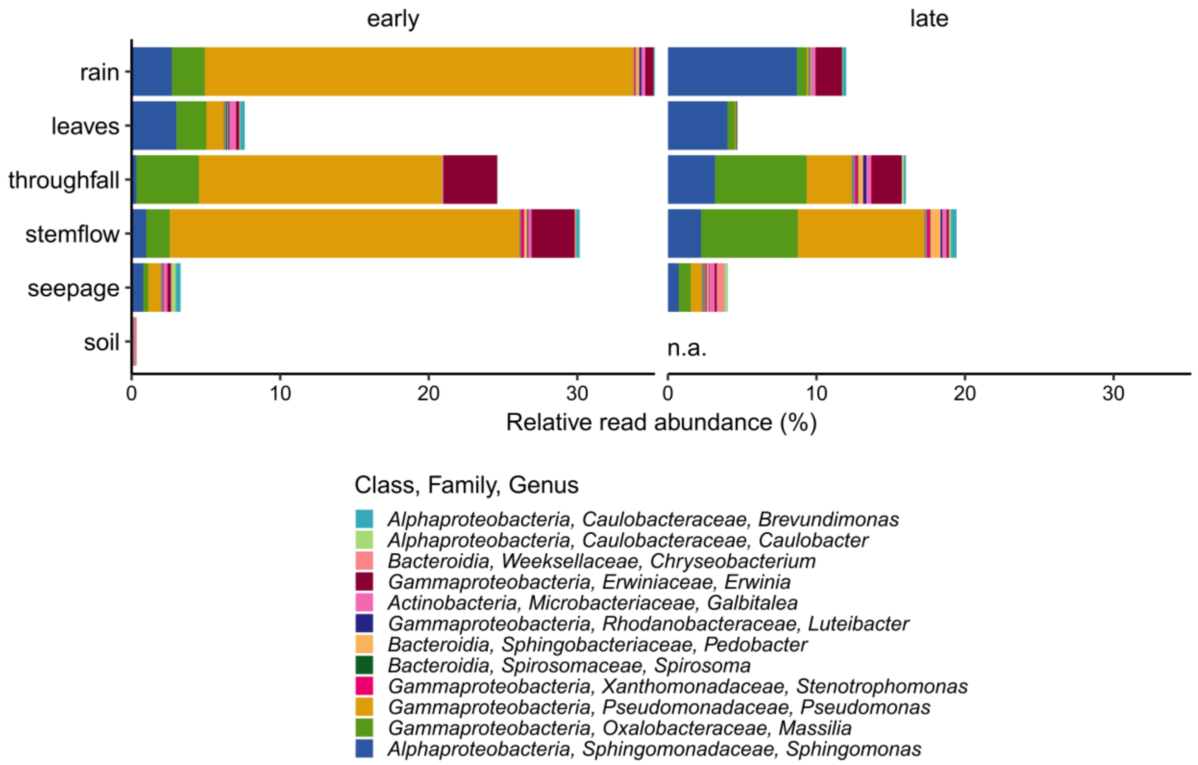


**Figure S 4:** Relative abundance of the 23 ASVs (summarized on genus level) which were shared across all six compartments and both sampling time points.

### Figure S5

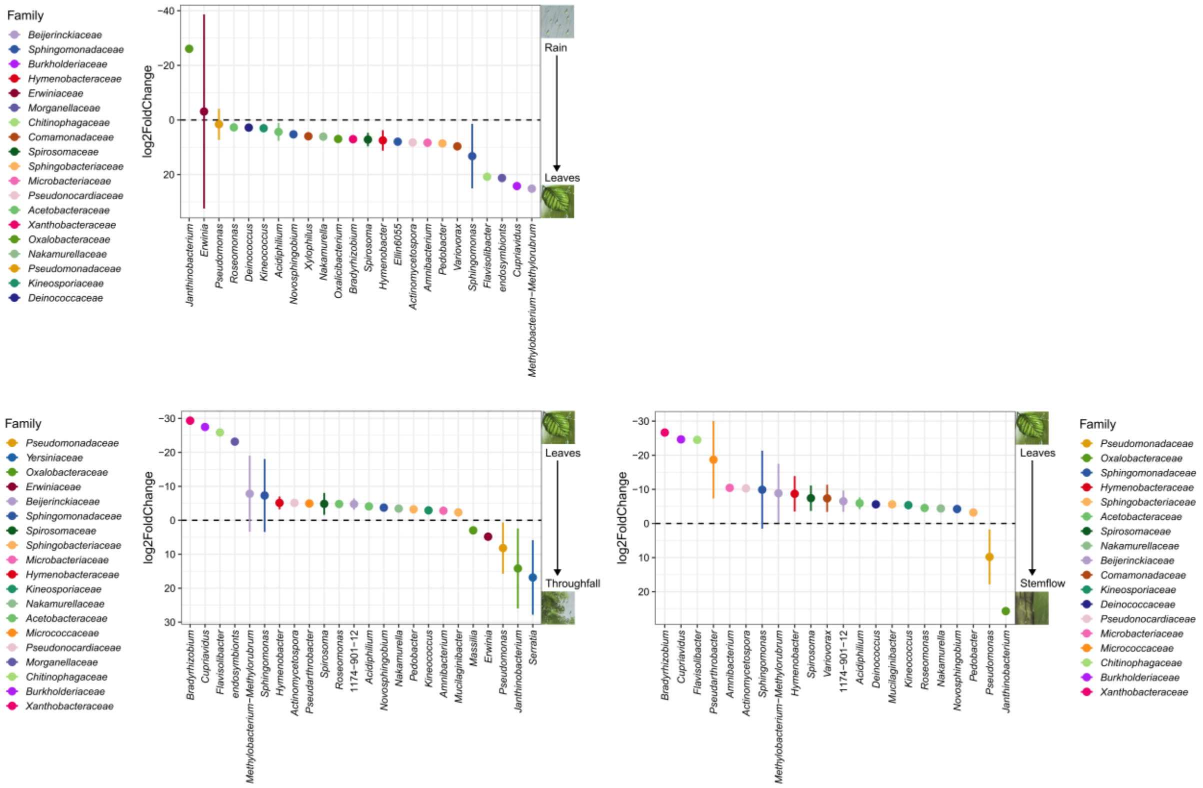


**Figure S 5:** Differential abundance analysis (Deseq2) of bacterial AVS (without singletons) on genus-level (color code summarized on family-level) showing the prevalence of being exported by rain to leaf surfaces and washed off from leaves by throughfall and stemflow.

### Figure S6

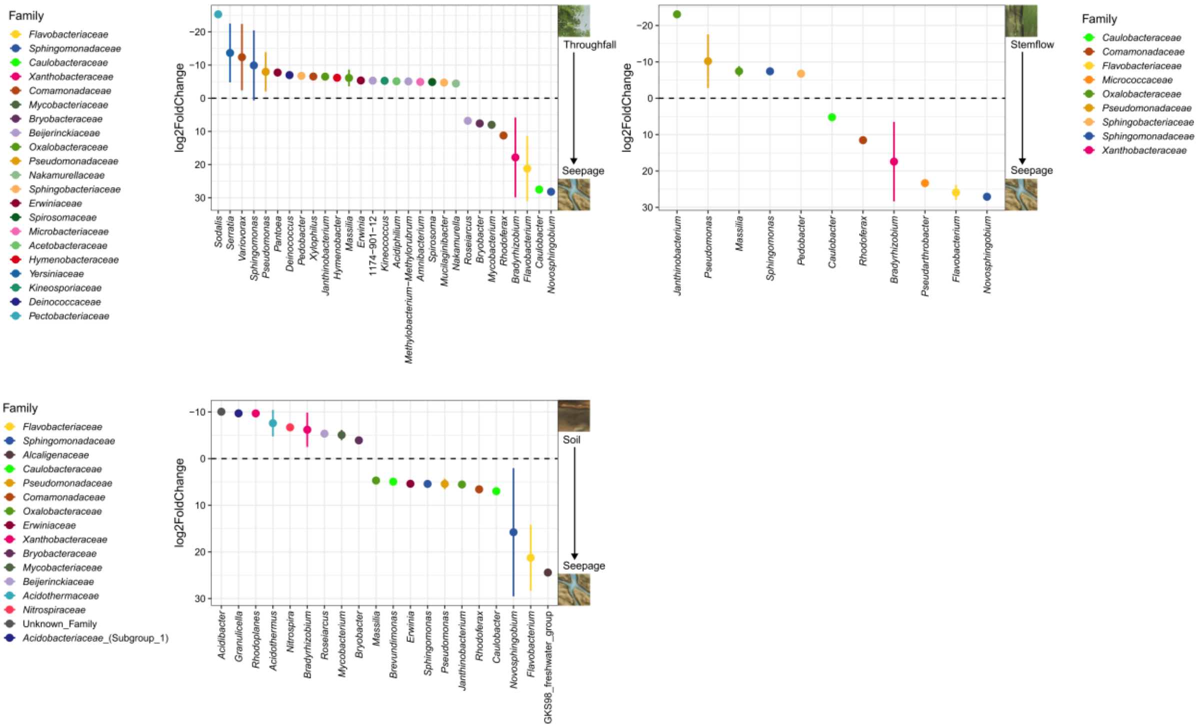


**Figure S 6:** Differential abundance analysis (Deseq2) of bacterial AVS (without singletons) on genus-level (color code summarized on family-level) showing the prevalence to be enriched in seepage from throughfall, stemflow and soil.

#### Figure S7

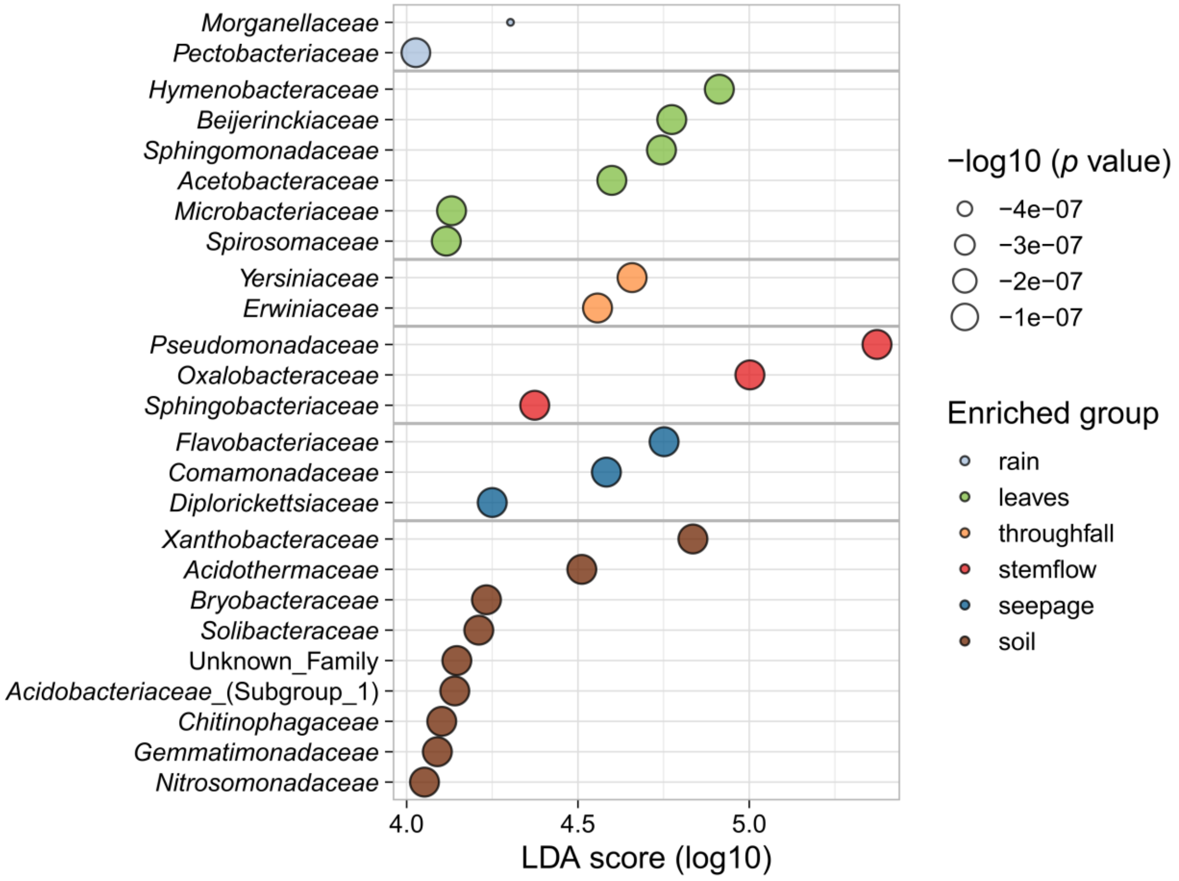


**Figure S 7:** Enrichment of most prevalent bacterial families in each compartment determined by LEFse. LDA value depicts strength of enrichment with *p* values displayed by the size.

#### Table S1

**Table S 1:** Overview of sampling volumes, microbial gene abundances $L^{-1}$/ $g^{-1}$ dw (gram dry weight), carbon and nitrogen concentrations from all compartments from May and September. EC = electrical conductivity, TOC = total organic carbon, POC = particulate organic carbon, DOC = dissolved organic carbon, IC = inorganic carbon, TN = total nitrogen, TPN = total particulate nitrogen, TDN = total dissolved nitrogen, DON = dissolved organic nitrogen, DIN = dissolved inorganic nitrogen. Values in brackets depict the standard deviation of the mean.

116

| Compartment | Growing season | Sampling Volume ($L^{-1}$) | Bacteria 16S rRNA ($u^{-1}$) | Estimated bacterial cells ($u^{-1}$) | Archaea 16S rRNA ($u^{-1}$) | Shannon (bacteria) | Richness (bacteria) | pH | EC | TOC (mg $L^{-1}$) | POC (mg $L^{-1}$) | DOC (mg $L^{-1}$) | IC (mg $L^{-1}$) | TN (mg $L^{-1}$) | TPN (mg $L^{-1}$) | TDN (mg $L^{-1}$) |
|---|---|---|---|---|---|---|---|---|---|---|---|---|---|---|---|---|
| **rain (u = L)** | early (n = 4) | 0.47 (0.01) | $2.79 \times 10^{8}$ ($2.37 \times 10^{8}$) | $7.20 \times 10^{7}$ ($5.79 \times 10^{7}$) | $2.10 \times 10^{5}$ ($1.21 \times 10^{5}$) | 3.46 (0.45) | 152.25 (67.83) | 6.67 (0.17) | 13.45 (0.99) | 6.06 (1.13) | 0.98 (1.03) | 5.08 (1.44) | 0.52 (0.04) | 1.57 (0.25) | 0.13 (0.04) | 1.44 (0.22) |
| | late (n= 4) | 0.45 (0.05) | $7.89 \times 10^{6}$ ($3.71 \times 10^{6}$) | $2.77 \times 10^{6}$ ($1.26 \times 10^{6}$) | $7.39 \times 10^{4}$ ($6.67 \times 10^{4}$) | 4.37 (0.21) | 235.25 (36.94) | 6.77 (0.63) | 2.13 (0.75) | 1.04 (0.23) | 0.08 (0.12) | 0.97 (0.11) | 0.05 (0.00) | 0.29 (0.04) | 0.02 (0.01) | 0.27 (0.03) |
| **leaves (u = $g^{-1}$ dw)** | early (n = 24) | - | $3.82 \times 10^{6}$ ($4.61 \times 10^{6}$) | $1.19 \times 10^{6}$ ($1.44 \times 10^{6}$) | $1.04 \times 10^{4}$ ($7.68 \times 10^{3}$) | 4.74 (0.71) | 321.54 (108.28) | - | - | - | - | - | - | - | - | - |
| | late (n = 24) | - | $6.53 \times 10^{8}$ ($5.43 \times 10^{8}$) | $2.04 \times 10^{8}$ ($1.70 \times 10^{8}$) | $3.20 \times 10^{4}$ ($2.91 \times 10^{4}$) | 4.76 (0.39) | 309.46 (54.75) | - | - | - | - | - | - | - | - | - |
| **throughfall (u = L)** | early (n = 5) | 0.34 (0.06) | $3.80 \times 10^{11}$ ($3.25 \times 10^{11}$) | $7.09 \times 10^{10}$ ($6.92 \times 10^{10}$) | $1.44 \times 10^{6}$ ($6.05 \times 10^{5}$) | 2.69 (0.51) | 116.60 (50.09) | 6.21 (0.29) | 57.69 (20.04) | 45.7 (32.12) | 14.32 (13.80) | 31.38 (19.29) | 3.60 (1.04) | 4.72 (1.75) | 2.25 (1.85) | 2.47 (0.80) |
| | late (n = 5) | 0.47 (0.07) | $3.08 \times 10^{10}$ ($2.23 \times 10^{10}$) | $8.72 \times 10^{9}$ ($6.13 \times 10^{9}$) | $1.22 \times 10^{6}$ ($1.16 \times 10^{6}$) | 4.70 (0.41) | 342.11 (69.18) | 6.40 (0.32) | 22.28 (13.30) | 12.91 (9.40) | 1.27 (0.81) | 11.64 (8.64) | 1.48 (0.81) | 1.33 (0.25) | 0.16 (0.08) | 1.16 (0.21) |
| **stemflow (u = L)** | early (n = 4) | 3.43 (2.63) | $3.12 \times 10^{11}$ ($2.94 \times 10^{11}$) | $9.07 \times 10^{10}$ ($9.62 \times 10^{10}$) | $8.47 \times 10^{5}$ ($4.39 \times 10^{5}$) | 3.14 (1.29) | 220.25 (145.17) | 6.47 (0.21) | 198.70 (96.77) | 190.45 (90.40) | 4.85 (3.72) | 185.60 (92.03) | 5.99 (3.20) | 13.14 (6.97) | 1.63 (1.27) | 11.51 (6.87) |
| | late (n = 4) | 8.92 (3.24) | $3.85 \times 10^{10}$ ($1.71 \times 10^{10}$) | $1.15 \times 10^{10}$ ($4.93 \times 10^{9}$) | $2.61 \times 10^{5}$ ($1.14 \times 10^{5}$) | 5.19 (0.45) | 580.75 (140.39) | 7.06 (0.08) | 78.00 (42.53) | 50.10 (31.50) | 1.66 (0.93) | 48.45 (30.68) | 4.17 (1.54) | 3.44 (1.62) | 0.14 (0.06) | 3.30 (1.64) |
| **seepage litter (u = L)** | early (n = 4) | 0.26 (0.07) | $5.44 \times 10^{10}$ ($2.36 \times 10^{10}$) | $1.7 \times 10^{10}$ ($7.38 \times 10^{10}$) | $1.49 \times 10^{7}$ ($9.23 \times 10^{6}$) | 4.88 (0.70) | 484.00 (147.89) | 6.72 (0.12) | 189.13 (46.48) | 70.07 (29.88) | 6.27 (4.22) | 63.80 (34.04) | 2.11 (0.57) | 13.92 (6.61) | 0.45 (0.36) | 13.46 (6.89) |
| | late (n = 4) | 0.77 (0.13) | $5.14 \times 10^{10}$ ($4.94 \times 10^{10}$) | $1.61 \times 10^{10}$ ($1.55 \times 10^{10}$) | $5.17 \times 10^{6}$) ($5.94 \times 10^{6}$) | 4.78 (0.48) | 349.75 (130.73) | 6.63 (0.21) | 197.00 (86.74) | 44.66 (8.52) | 1.31 0.64) | 63.80 8.04) | 3.73 (3.22) | 17.60 11.83) | 0.73 0.99) | 16.88 10.90) |
| **seepage 0-4 cm (u = L)** | early (n = 2) | 0.26 (0.10) | $2.80 \times 10^{10}$ ($1.84 \times 10^{10}$) | $8.75 \times 10^{9}$ ($5.74 \times 10^{9}$) | $2.84 \times 10^{8}$ ($3.74 \times 10^{8}$) | 4.15 (0.03) | 245.00 (120.21) | 6.74 (0.09) | 145.95 (80.68) | 43.47 (22.64) | 3.58 (0.48) | 39.89 (23.12) | 2.11 (2.42) | 11.44 (6.44) | 0.18 (0.21) | 11.26 (6.64) |
| | late (n = 4) | 0.67 (0.33) | $2.75 \times 10^{10}$ ($2.13 \times 10^{10}$) | $8.59 \times 10^{9}$ ($6.64 \times 10^{9}$) | $1.69 \times 10^{7}$ ($3.21 \times 10^{7}$) | 4.69 (0.25) | 257.75 (55.68) | 6.32 (0.20) | 73.50 (19.26) | 30.99 (7.88) | 1.45 (0.35) | 29.53 (7.64) | 2.98 (0.83) | 3.89 (2.09) | 0.12 (0.07) | 3.77 (2.10) |
| **seepage 0-16 cm (u = L)** | early (n = 2) | 0.34 (0.10) | $6.27 \times 10^{9}$ ($1.07 \times 10^{9}$) | $1.96 \times 10^{9}$ ($3.35 \times 10^{8}$) | $7.34 \times 10^{8}$ ($9.37 \times 10^{8}$) | 5.39 (0.74) | 498.00 (292.74) | 5.36 (0.18) | 150.20 (6.22) | 12.25 (6.71) | 0.06 (0.08) | 12.18 (6.78) | 0.23 (0.01) | 13.29 (1.72) | 0.04 (0.01) | 13.25 (1.70) |
| | late (n = 4) | 0.48 (0.38) | $1.30 \times 10^{10}$ ($1.36 \times 10^{10}$) | $4.08 \times 10^{9}$ ($4.26 \times 10^{9}$) | $3.14 \times 10^{8}$ ($5.15 \times 10^{8}$) | 5.26 (0.39) | 424.25 (197.48) | 5.74 (0.81) | 201.50 (119.56) | 29.27 (29.88) | 1.09 (1.18) | 22.43 (26.12) | 1.02 (1.74) | 20.65 (16.89) | 0.29 (0.31) | 20.44 (16.76) |
| **seepage 0-30 cm (u = L)** | early (n = 3) | 0.52 (0.20) | $3.43 \times 10^{9}$ ($2.02 \times 10^{9}$) | $1.07 \times 10^{9}$ ($6.32 \times 10^{8}$) | $9.70 \times 10^{8}$ ($6.33 \times 10^{8}$) | 5.82 (0.25) | 649.50 (17.68) | 6.09 (0) | 133.80 (10.47) | 8.16 (5.95) | 0.43 (0.36) | 7.73 (5.62) | 11.38 (1.80) | 13.84 (5.55) | 0.50 (0.84) | 13.34 (4.72) |
| | late (n = 4) | 0.55 (0.30) | $1.03 \times 10^{10}$ ($7.04 \times 10^{9}$) | $3.22 \times 10^{9}$ ($2.20 \times 10^{9}$) | $1.23 \times 10^{8}$ ($1.10 \times 10^{8}$) | 5.33 (0.25) | 434.50 (125.53) | 6.64 (0.88) | 207.50 (99.66) | 7.48 (7.69) | 0.52 (0.70) | 8.82 (6.82) | 4.76 (8.49) | 18.04 (12.65) | 0.28 (0.18) | 17.82 (12.61) |
| **soil 0-4 cm (u = gdw)** | early (n = 12) | - | $1.62 \times 10^{10}$ ($4.77 \times 10^{9}$) | $5.05 \times 10^{9}$ ($1.49 \times 10^{9}$) | $3.36 \times 10^{7}$ ($2.17 \times 10^{7}$) | 5.57 (0.44) | 480.08 (169.51) | 5.49 (0.54) | - | - | - | - | - | - | - | - |
| **soil 4-16 cm (u = gdw)** | early (n = 12) | - | $1.44 \times 10^{10}$ ($5.87 \times 10^{9}$) | $4.49 \times 10^{9}$ ($1.84 \times 10^{9}$) | $2.41 \times 10^{7}$ ($1.75 \times 10^{7}$) | 5.38 (0.39) | 395.18 (138.24) | 5.23 (0.40) | - | - | - | - | - | - | - | - |
| **soil 16-30 cm (u = gdw)** | early (n = 11) | - | $8.41 \times 10^{9}$ ($3.99 \times 10^{9}$) | $2.63 \times 10^{9}$ ($1.25 \times 10^{9}$) | $3.39 \times 10^{7}$ ($2.14 \times 10^{7}$) | 5.01 (0.30) | 288.45 (93.42) | 5.18 (0.36) | - | - | - | - | - | - | - | - |
